# Granular Extracellular Matrix (gECM) Hydrogel Wafers as Shelf-Stable 2.5D Substrates for Microphysiological Modeling

**DOI:** 10.64898/2026.05.26.727948

**Authors:** Juliet O. Heye, Shannon A. Blanco, Stephanie E. Schneider, Shantae Gallegos, Jeanne E. Barthold, Michael Floren, Victor Acosta, Stephane Avril, Corey P. Neu

## Abstract

Understanding disease pathology and evaluating emerging therapeutics require *in vitro* models that accurately recapitulate human tissue environments. However, existing microphysiological systems often compromise either biomimicry or ease-of-use, limiting widespread adoption and scalability. Here, we present lyophilized granular extracellular matrix (gECM) hydrogel wafers as shelf-stable, humanized 2.5D substrates that enable physiologically relevant modeling while simplifying integration into experimental workflows. Derived from decellularized human cartilage and bone, gECM hydrogel wafers retain tissue-specific architecture while introducing microporosity and surface topography through lyophilization. These wafers maintain swelling behavior, structural integrity, and mechanical properties over three months of room-temperature storage, allowing pre-fabrication and on-demand use without loss of function. gECM hydrogel wafers support direct cell seeding without encapsulation and sustain viability and proliferation of human adipose-derived mesenchymal stromal cells over 21 days, with gene expression trends comparable to 3D gECM hydrogels. Furthermore, wafers can be readily integrated into microfluidic systems with *in situ* hydration and transport of large biomolecules. Together, this platform bridges the gap between conventional 2D culture ease-of-use and 3D biomaterial biomimicry, providing a scalable and physiologically relevant *in vitro* model approach for high-throughput disease studies and therapeutic screening.

## INTRODUCTION

Microphysiological models that recapitulate native tissue are critical for enabling *in vitro* investigation of disease pathology and emerging therapeutics [1–3]. Recent regulatory and funding changes, including the FDA Modernization Act 2.0 [4,5] and increased NIH emphasis on New Approach Methods [6], have pushed a shift away from animal models toward human-relevant *in vitro* systems [7–11]. These changes are driven in part by the high failure rate of therapeutics in clinical trials [12], which is often attributed to the limited predictive power of pre-clinical models and the inability of animal systems to fully capture human physiology [1–3]. However, current *in vitro* technologies remain insufficient to accurately represent native tissue environments, hindering the lab-to-clinic translation of new therapeutics. This gap highlights the need for humanized, tissue-specific substrates that are both biologically relevant and broadly accessible for pre-clinical testing. One disease that underscores this limitation is osteoarthritis, a highly prevalent and poorly understood condition for which no disease-modifying therapies currently exist [13]. Although traditionally defined as a cartilage disease [13], osteoarthritis is now recognized as a multi-tissue pathology involving cartilage, subchondral bone, and surrounding joint tissues [14,15]. This complexity necessitates model systems capable of recapitulating multiple tissue environments and their interactions, enabling more accurate study of disease progression and therapeutic response [10,11,16].

A central challenge in developing biomimetic tissue models is balancing physiological relevance with usability. Conventional 2D culture systems are widely used due to their simplicity, but they lack appropriate mechanical and structural cues, resulting in altered cell behavior [17–21]. For example, chondrocytes cultured in 2D have been shown to dedifferentiate over time and exhibit mechanical memory, limiting their ability to recover tissue-specific phenotypes even after transfer to more biomimetic environments [18]. While coatings such as Matrigel are commonly used to introduce biological signals, they are derived from mouse tumor tissue and lack tissue-specific relevance [20,21]. In contrast, 3D scaffold-based systems provide improved structural cues, but require cell encapsulation, polymerization, or specialized fabrication workflows that limit accessibility and introduce potential for unwanted variability [22–25]. Within 3D systems, bulk hydrogels—whether synthetic or naturally derived—are widely used due to their tunability. However, they often require trade-offs between mechanical integrity and biological relevance. Synthetic hydrogels offer robust and controllable mechanical properties but lack native biochemical signals, while naturally derived hydrogels provide improved biological cues but frequently exhibit weak mechanical properties and limited structural stability. Microgel-based systems have been developed to improve mechanical performance and injectability [26–29], but these materials still lack the tissue-specific microarchitecture characteristic of native extracellular matrix.

Extracellular matrix (ECM)-derived biomaterials offer an alternative strategy by preserving native tissue composition [30]. However, intact ECM tissues or grafts are difficult to source and control in size and geometry [31,32], while digestion into soluble ECM compromises mechanical integrity and eliminates native microstructure [33,34]. Granular ECM (gECM) hydrogels address these limitations by processing tissues into particulate form, preserving both compositional and micro-architectural cues, and densely packing them into a hydrogel base to form solidified constructs [22,23,35]. Unlike conventional hydrogels, gECM materials uniquely couple tissue-specific composition with preserved microstructure, enabling tunable mechanical properties governed by particle packing and percolation [23]. Prior work with a thiolated hyaluronic acid (tHA) -based gECM hydrogel demonstrated that tissue type drives mechanics and promotes tissue-specific cellular responses [22,23]. However, these materials are typically implemented as 3D encapsulation platforms, which still require biomaterials expertise and limit accessibility.

To bridge the gap between physiological relevance and usability, 2.5D substrates have emerged as an intermediate approach that combines advantages of both 2D and 3D systems [36–42]. These platforms allow cells to be introduced directly onto a biomimetic substrate without encapsulation, while still providing structural and mechanical cues that influence cell behavior [43–47]. Substrate microstructure, including surface roughness and porosity, has been shown to influence lineage-specific cell activity [37,38,48–51], highlighting the importance of material architecture in directing cell fate. Fabrication strategies such as freezing and lyophilization introduce microporosity that promotes cell integration and facilitates transport of nutrients, signaling molecules, and therapeutics [46,52]. Importantly, lyophilization also enables room-temperature stability [53], supporting storage, transport, and high-throughput use. However, the stability of gECM hydrogel substrates in this form remains largely unexplored.

Biomimicry, ease of use, and shelf-stability are particularly advantageous for integration into microfluidic tissue-on-chip models. On-chip systems enable implementation of tissue-relevant mechanics, controlled delivery of nutrients and targeted therapeutics, and connection of multiple chips to create a multi-tissue system [10,11,54,55]. However, incorporation of tissue-specific biomaterials into these platforms remains challenging, as chip fabrication processes often require elevated temperatures or surface treatments that are incompatible with many biomaterials. As a result, many systems rely on cells flowed into pre-assembled chips alone, limiting their ability to fully recapitulate native tissue structure. Pre-formed, stable biomaterial substrates that can be readily integrated without additional processing would address this limitation and improve the physiological relevance of these models.

In this study, we fabricate lyophilized tHA-based gECM hydrogel wafers derived from human cartilage and bone as shelf-stable 2.5D substrates for microphysiological modeling. We characterize their biophysical and mechanical properties, including changes over a three-month room-temperature storage period to assess shelf stability, and evaluate microstructure and surface topology relevant to cell–material interactions. We further establish a simple cell seeding approach using human adipose-derived mesenchymal stromal cells (adMSCs) and assess cell viability, phenotypic response, and substrate stability in culture. Finally, we demonstrate integration of cartilage gECM wafers into a proof-of-concept microfluidic chip system, including *in situ* hydration and nutrient flow. Together, this work presents a scalable and user-friendly strategy for generating humanized, tissue-specific substrates that support physiologically relevant *in vitro* models for disease study and therapeutic screening.

## RESULTS

### Fabrication of human cartilage and bone gECM hydrogel wafers

We previously developed a decellularization protocol for porcine tissues [manuscript in progress] using an acid-base viral inactivation treatment [56], followed by post-decellularization particularization to form ECM particles with controlled particle size and reduced DNA content. We adapted this decellularization protocol and post-decellularization particularization to human cartilage and bone tissues that would otherwise be discarded from a human donor tissue bank (**Figure 1A**). We now use these decellularized human cartilage and bone particles to fabricate gECM hydrogel wafers (**Figure 1B**). We follow previously established formulations in combining cartilage and bone particles with thiolated hyaluronic acid (tHA) to form gECM hydrogels. The mixed gECM hydrogels are molded to a desired geometry, incubated at 37°C following previously established polymerization times, isotropically frozen at -20°C, and lyophilized to form gECM hydrogel wafers. The gECM hydrogel wafers appear white in color with noticeable porosity and surface texture (**Figure 1C**) and are stored dry at room temperature until use. Wafers are rehydrated in DPBS prior to characterization, appearing smoother and more swollen compared to their dry form.

**Figure 1.**
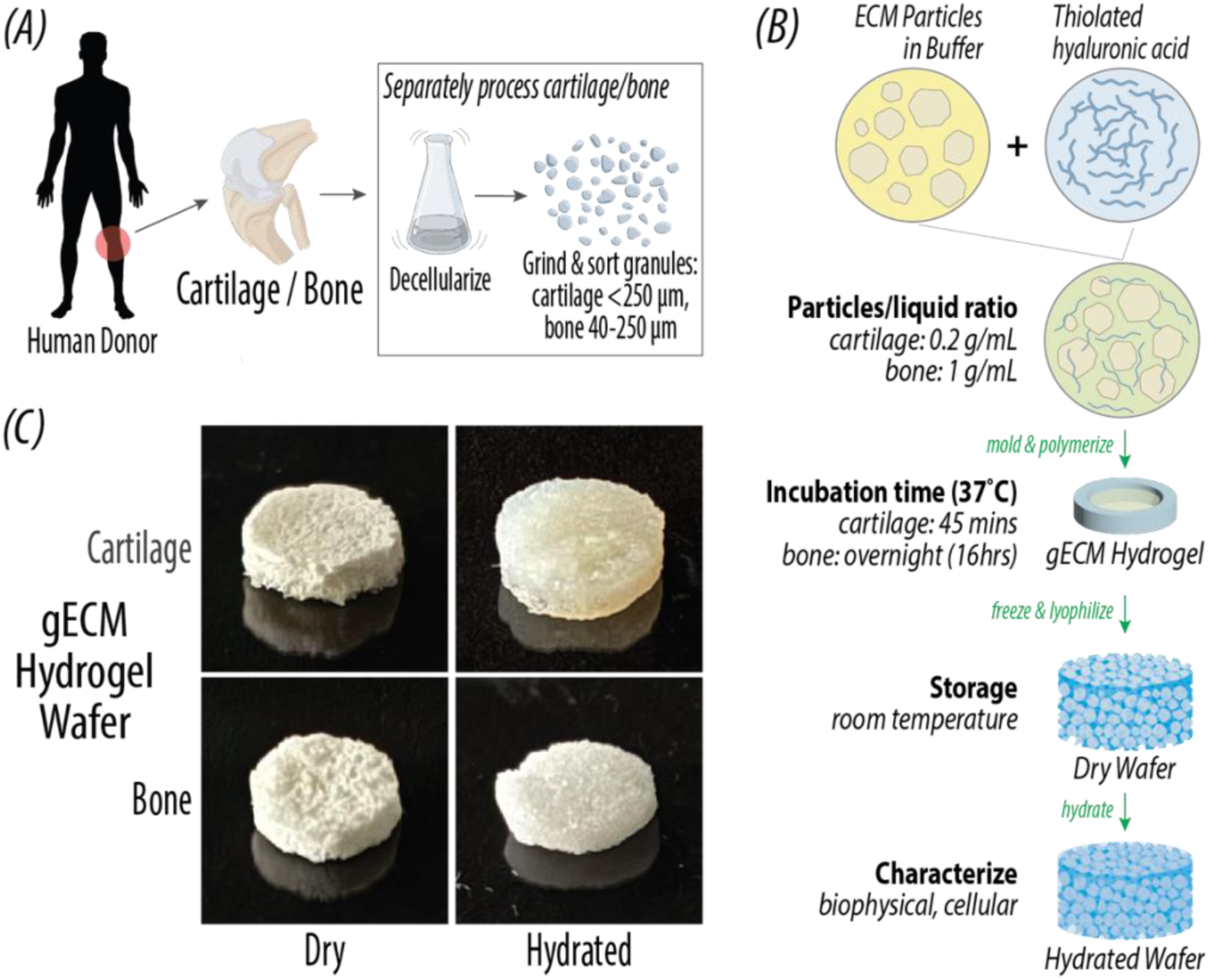
Fabrication of human cartilage and bone gECM hydrogel wafers. (A) Human cartilage and bone tissues are decellularized and size sorted. (B) ECM particles are combined with thiolated hyaluronic acid (tHA) to form gECM hydrogels. Polymerized gECM hydrogels are then isotropically frozen and lyophilized to form gECM hydrogel wafers. Wafers are rehydrated for characterization (DPBS >24hrs). (C) Macroscopic images of cartilage and bone gECM hydrogel wafers in dry and hydrated form.

### Room temperature storage does not affect swelling behavior of gECM hydrogel wafers

Application of gECM hydrogel wafers involves aqueous environments, so we measured the mass and geometry while submerged in DPBS to characterize rehydration behavior. We measured swelling behavior of gECM wafers immediately after fabrication (fresh), after 1 month of room temperature dry storage, and after 3 months of room temperature dry storage. Mass, volume, surface area, and height were measured for wafers in dry form and at 6 time points while swelling in DPBS (30 mins, 1hr, overnight ∼16hrs, 24hrs, 48hrs, 1 week). Values were normalized to the wafers when dry to quantify changes relative to their initial form (**Figure 2A**). Overall, we observe much greater capacity for swelling in cartilage compared to bone wafers. This is supported statistically, with significantly greater mass, volume, surface area, and height in cartilage compared to bone wafers at all time points past dry (all p<0.001). We see minimal differences in swelling patterns between fresh, 1-month, and 3-month wafers for both cartilage and bone wafers. Importantly, there was no statistical effect of shelf life on any swelling results, supporting that the water-uptake capacity of gECM wafers is stable over time. Cartilage wafers significantly increased in mass between dry and 30 mins, and again between 1hr and overnight time points, indicating that cartilage does not plateau in mass until after 24hrs of swelling. Bone significantly increased in mass only between dry and 30mins, supporting that it plateaus in mass after only 30mins of swelling.

**Figure 2.**
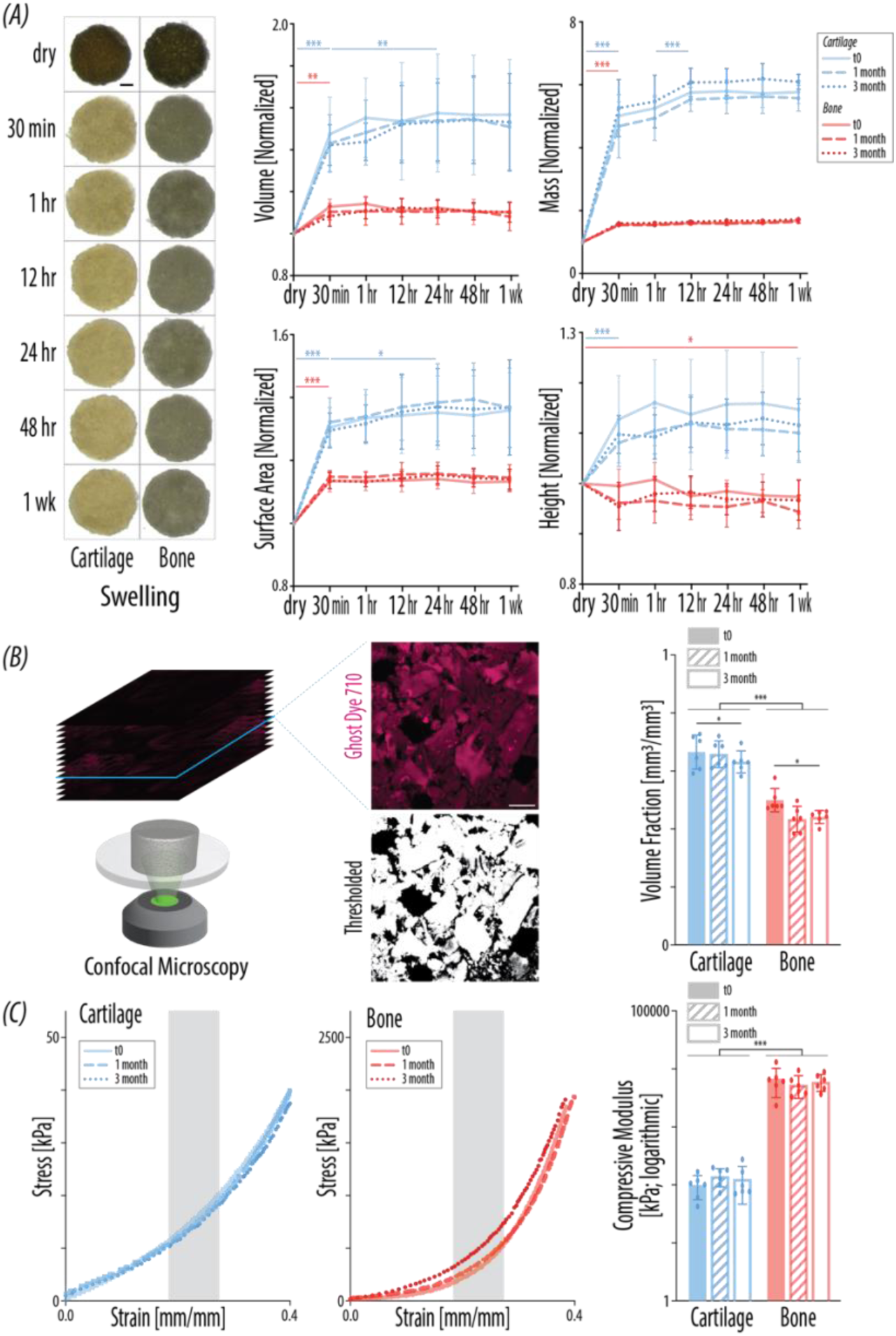
gECM hydrogel wafers maintain biophysical and mechanical characteristics over a 3-month shelf-life study. All characterization was performed on gECM hydrogel wafers immediately after fabrication (fresh), after 1 month of dry room-temperature storage, and after 3 months of dry room-temperature storage. (A) Representative top-down images of gECM hydrogel wafers during swelling, used to quantify surface area. Scale bar = 1mm. Mass, volume, surface area, and height of gECM hydrogel wafers over the course of 1 week swelling in DPBS, normalized to initial dry form (N = 6). (B) Representative confocal slices through optical depth of fully swelled cartilage gECM hydrogel wafer. Selected slices were thresholded in MATLAB, and area fraction of particles was averaged across slices to calculate volume fraction of particles to total volume (N = 6). Scale bar = 100 µm. (C) Representative stress-strain curves of gECM hydrogel wafers during bulk unconfined compression. Bulk compressive moduli of gECM hydrogel wafers calculated between 20-30% strain (N=6). For all plots, error bars = standard deviation, *p<0.05, **p<0.01, ***p<0.001.

For volume, cartilage wafers significantly increased between dry and 30min time points and again between 30mins and 24hrs, suggesting that cartilage volume also plateaus after 24hrs. Similar to mass, bone wafer volume plateaued after only 30 mins, with a significant increase in volume only between dry and 30 mins. We further characterized the surface area and height during swelling to understand whether gECM wafers swell uniformly or not. This provides precise dimensional control of gECM wafers, facilitating their integration into *in vitro* systems, including microfluidic platforms where geometric constraints are critical. Cartilage significantly increased in surface area between dry and 30 mins and between 30 mins to 24 hrs, but only significantly increased in height between dry and 30 mins. With a ∼1.5-fold increase in volume, ∼1.3-fold increase in surface area, and ∼1.1-fold increase in height, we can conclude that cartilage swelled uniformly in all axes. However, the height reached its max height earlier than surface area. In contrast, bone significantly increased in surface area between only dry and 30 mins, but significantly decreased in height between dry and 1 week. This suggests that the ∼1.1-fold increase in bone wafer volume is primarily drive by expansion laterally, with possibly some compaction in the z-axis. This may be a result of the heavier mineralized bone particles settling with time. Patterns in height are additionally confirmed by optical coherence tomography (OCT) images, which show minimal changes in height in bone wafers and increase in height in cartilage wafers between dry and 24hr hydrated states (**Supplementary Figure 1A**). We additionally show raw swelling measurements (**Supplementary Figure 1B**), which shows that while bone swells less overall with lower volume compared to cartilage, its high mineral content results in higher mass compared to cartilage. Overall, cartilage plateaus in swelling by 24hrs, while bone modestly swells in 30 mins and after that stays effectively constant over 1 week.

### gECM hydrogel wafers maintain mechanics despite modest decrease in particle volume fraction during room temperature storage

We then quantified particle-to-total volume fraction for fully-swelled gECM hydrogel wafers (**Figure 2B**). Overall, cartilage had a significantly higher particle volume fraction compared to bone wafers. Interestingly, the particle volume fraction for cartilage gECM hydrogel wafers (∼0.65) is notably higher than previously measured for cartilage gECM hydrogels (∼0.55). In contrast, bone gECM hydrogel wafers have modestly higher volume fraction (∼0.50) compared to previously measured bone gECM hydrogels (∼0.45) when characterized fresh, and comparable volume fraction (∼0.44) at 1 and 3 months. The high pressure induced during lyophilization likely compacts particles together, but it appears that the denser mineralized bone particles may resist this compaction compared to cartilage. Statistically, we see a significant but modest overall decrease in volume fraction between fresh and 3-month wafers for both cartilage and bone gECM hydrogel wafers. We do observe a more noticeable decrease in volume fraction for bone between fresh and later shelf-life time points, suggesting that bone does not maintain the higher compaction state over time as well as cartilage. Importantly, this observed lower packing density in bone appears to remain steady between 1 month and 3 months, suggesting that it is settled at the pre-lyophilization volume fraction, potentially maintained by the tHA network.

We then measured bulk stiffness of fully swelled gECM hydrogel wafers. Representative stress-strain curves revealed a J-shape for all wafers, with lower stress in cartilage compared to bone wafers and minimal differences between shelf-life time points (**Figure 2C**). We calculated the compressive moduli between 20-30% strain where the effect of percolation is exaggerated. Additionally, some bone samples reached maximum load cell force in the 30-40% strain range. Compressive modulus calculated between 20-30% strain was significantly higher in bone compared to cartilage gECM hydrogel wafers, as expected. We saw this same trend across 4 regions of strain (0-10%, 10-20%, 20-30%, and 30-40%) (**Supplementary Figure 2**). We observe a slight decrease in compressive modulus (20-30% strain) between fresh and 1-month bone wafers, which aligns with the observed decrease in particle volume fraction. This matches previous work in which decreased particle volume fraction drives decreased bulk stiffness through percolation. Importantly however, there are no statistical differences in the compressive moduli of fresh, 1-month, or 3-month wafers for both cartilage and bone, despite modest decreases in particle volume fraction. Overall, the mechanics of gECM hydrogel wafers is stable over a 3-month room temperature storage period.

### gECM hydrogel wafers display defined internal microporosity and surface topography

Scanning electron microscopy (SEM) was used to characterize the microstructure of cartilage and bone gECM hydrogel wafers in dry and hydrated form (**Figure 3A**) and to compare these structures to non-lyophilized gECM hydrogels (**Supplementary Figure 3**). Compared to hydrogels, wafers exhibited notably increased porosity. This was particularly evident at the top surface and most pronounced in bone-derived materials. In hydrogels, the thiolated hyaluronic acid (tHA) phase appeared to form a relatively smooth layer over embedded ECM particles, limiting surface topographical features. In contrast, wafers displayed exposed ECM particles and pronounced interparticle void space. Within the internal structure, the tHA in hydrogels appeared more diffuse and fibrous, lacking the defined interconnected porosity observed in wafers.

**Figure 3.**
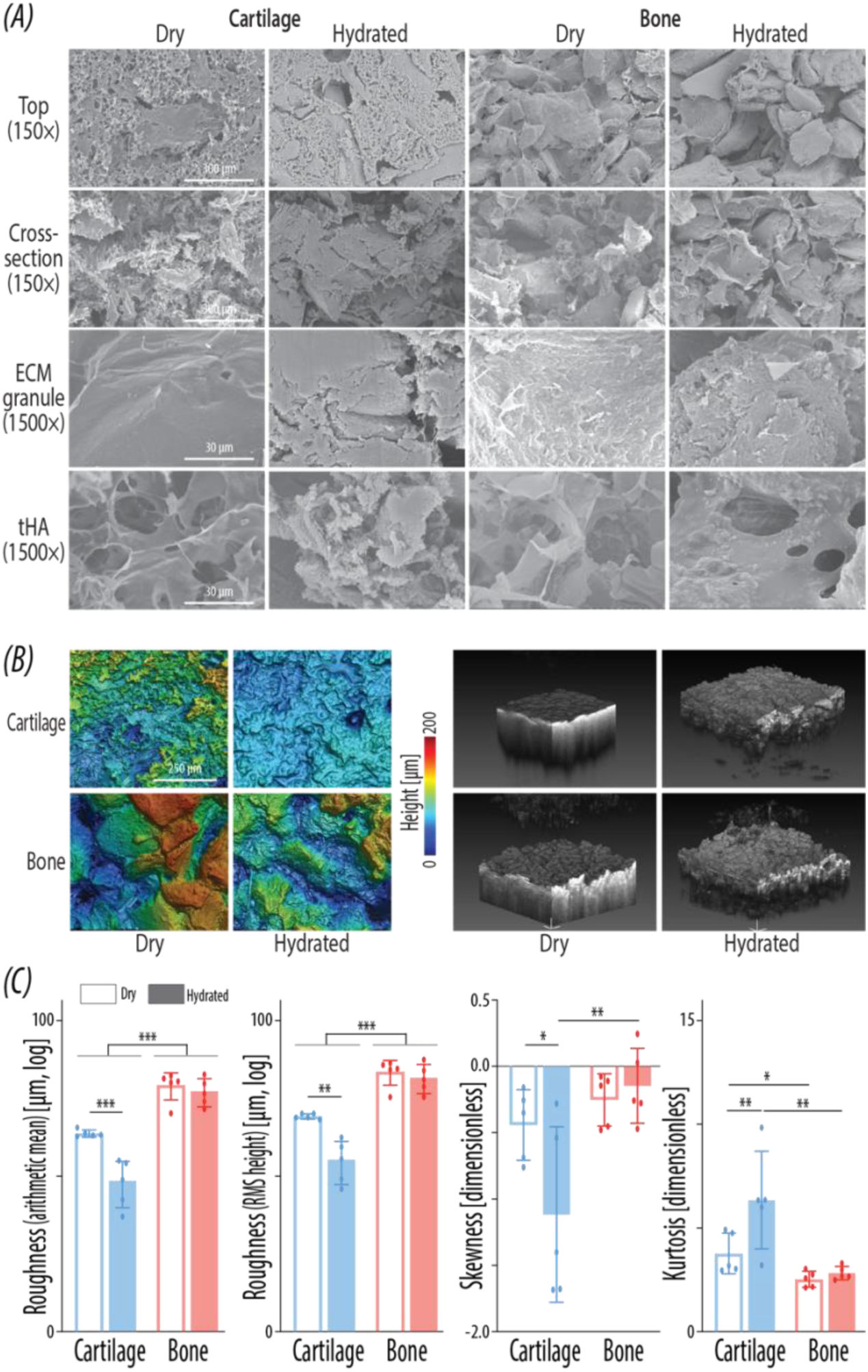
gECM hydrogel wafers display internal microporosity and surface topography. (A) Representative SEM images of cartilage and bone gECM hydrogel wafers in dry and hydrated (DPBS >24hrs) form. Images of the top and cross-section are at 150X magnification (scale bars = 300µm). Image of the particle and tHA are at 1500X magnification (scale bars = 30µm). (B) Left: Representative 3D laser scanning displays of surface roughness on cartilage and bone gECM hydrogel wafers in dry and hydrated (DPBS >24hrs) form. Color bar indicates height (0-200µm) across all images. Scale bar = 250µm. Right: Representative optical coherence tomography (OCT) images showing top surface of cartilage and bone gECM wafers in dry and hydrated (24hrs PBS) form. (C) Quantified surface roughness parameters for cartilage and bone gECM hydrogel wafers in dry and hydrated form (N = 5). For all plots, error bars = standard deviation, *p<0.05, **p<0.01, ***p<0.001.

Hydration of gECM hydrogel wafers resulted in observable changes in both surface and internal microstructure. Dry wafers exhibited sharper, more defined tHA structures with increased apparent microporosity, and ECM particles appeared more angular and rougher. Upon hydration, both the tHA network and ECM particle surfaces appeared smoother, with a slight reduction in microporosity. However, the overall porous architecture remained intact. While microporosity overall appears more distinct in dry compared to hydrated wafers, some larger deeper pores are observed in hydrated wafers. Differences between dry and hydrated states were more pronounced in cartilage than in bone, especially at the top surface. We also see distinct differences in surface and internal particle structure between cartilage and bone wafers. The bone particles are visually distinct from the surrounding matrix, while the cartilage particles appeared more integrated within the tHA network. Overall, SEM imaging reveals that gECM hydrogel wafers exhibited more defined interconnected porosity than gECM hydrogels, indicating that lyophilization transforms gECM hydrogels into a more porous and structurally accessible scaffold. Dry wafers displayed sharper and more defined pore structures, while hydrated wafers display smoother features. These structural characteristics may facilitate cellular integration and transport within the material.

Surface topology is qualitatively shown in 3D laser scanning surface displays and optical coherence tomography (OCT) images (**Figure 3B**). Surface roughness analysis revealed differences in both the magnitude and distribution of surface features, with arithmetic mean height and root mean square height describing overall roughness, while surface skewness and surface kurtosis provided insight into the asymmetry and extremity of surface topography (**Figure 3C**). Arithmetic mean height was significantly higher in bone compared to cartilage across conditions, indicating a generally rougher surface in bone-derived wafers. Hydration resulted in a significant decrease in arithmetic mean height for cartilage, while bone exhibited a slight but non-significant decrease. These trends were consistent with SEM observations, where cartilage surfaces appeared smoother following hydration, whereas bone retained more pronounced surface features. Comparable values and trends were observed in root mean square height, suggesting that the overall trends in roughness were not driven by extreme outlier features, as root mean square height is more sensitive to sharp peaks and deep pits.

Surface skewness values were negative across all conditions, indicating that surface topography was dominated by pits rather than peaks, consistent with the presence of surface-connected porosity. Hydrated bone exhibited significantly higher (less negative) skewness compared to hydrated cartilage, with values closer to zero, suggesting a more balanced distribution of peaks and pits. In contrast, hydrated cartilage exhibited significantly more negative skewness compared to its dry counterpart, indicating that surface roughness became increasingly dominated by valleys following hydration. Hydrated cartilage also exhibited significantly higher surface kurtosis compared to dry cartilage. Higher skewness and kurtosis together indicate the presence of more localized or extreme surface valleys in hydrated cartilage wafers. These features are consistent with the enhancement of discrete macropores, as observed in SEM, 3D surface maps, and OCT images. Additionally, kurtosis was significantly higher in cartilage compared to bone under both dry and hydrated conditions, suggesting that cartilage surfaces contain more pronounced localized features, whereas bone surfaces are comparatively smoother with broader, more rounded topographical features. Overall, these results indicate that hydration results in a generally smoother surface but with increased prominence of localized pits in cartilage wafers, while bone wafers maintain a rougher overall surface with more rounded and evenly distributed features. Across both tissue types, surface roughness is primarily driven by porosity. These distinct, tissue-dependent surface characteristics are likely to influence cell–material interactions and may contribute to differential cellular responses on gECM wafers.

### Pre-hydrated gECM hydrogel wafers support adMSC viability during seeding

Before assessing gECM hydrogel wafer cytocompatibility in long-term culture, we first compared different seeding methods to optimize human adMSC viability and integration into wafers. We seeded adMSCs suspended in culture medium on dry and pre-hydrated gECM hydrogel wafers (**Figure 4A**). Pre-hydrated wafers were submerged in culture medium for 24-48hrs and then medium was removed prior to seeding. Two hours after seeding adMSCs on both dry and hydrated cartilage and bone wafers, culture medium was added to cell-laden wafers. LIVE/DEAD imaging was performed on Day 3 of culture, looking at the top, cross-sectional, and bottom faces of the wafers (**Figure 4B**). Overall, we see greater viability in cells seeded on pre-hydrated wafers compared to cells seeded on dry wafers, particularly on the top surface. Cross-sectional images show that cells penetrated deeper into the dry wafers compared to hydrated wafers. Cells span the entire depth of dry wafers captured in images (∼1.1mm, total depth of wafers ∼1.5mm), with evidence of full depth penetration in the images of the bottom surface. However, a large majority of the cells appear dead. In contrast, the cells in the cross-sectional images of pre-hydrated wafers have better viability, but the viable cells only penetrate ∼250-500µm into the wafer. Some cells are observed deeper than that, but they appear to be dead. Cell death may be increased in dry wafers due to less available media during attachment time or due to contact with sharper features in the dry wafers observed in SEM imaging. It is also possible that dead cells were able to fall deeper into wafers due to their smaller diameter (∼8-10µm) compared to live cells (∼15-17µm). Taken together, these results provided clear indication to move forward with pre-hydrated wafers in subsequent cell studies.

**Figure 4.**
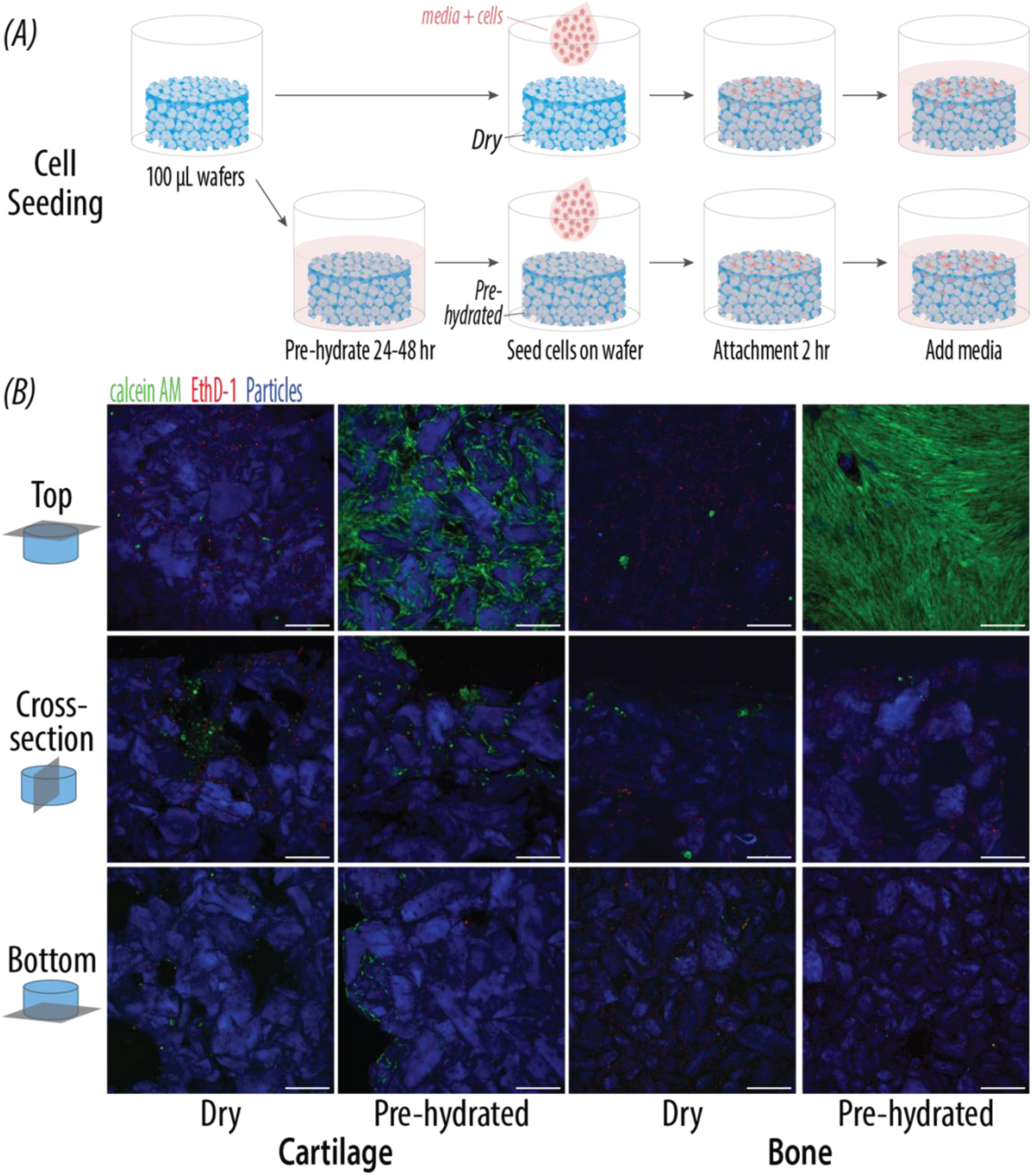
Pre-hydrated gECM hydrogel wafers support adMSC viability during seeding. (A) Ad-MSC’s were seeded onto dry and pre-hydrated cartilage and bone gECM hydrogel wafers. (B) Confocal images of the top, cross-section, and bottom faces of cartilage and bone gECM hydrogel wafers at Day 3. For all images, green = LIVE, red = DEAD, blue = ECM particles autofluorescence, Scale bars = 250µm.

In our seeding method study, we seeded 500,000 cells/100µL wafer to align with past work encapsulating cells into gECM hydrogels [22,23]. Here, we observed very high confluency of adMSCs on the top surface of pre-hydrated wafers at only Day 3 in culture. Therefore, we conducted an additional study to optimize wafer seeding density. Human adMSCs were seeded onto pre-hydrated cartilage and bone gECM hydrogel wafers at 250,000 cells, 125,000 cells, and 70,000 cells per 100µL wafer (**Supplementary Figure 4**). At Day 3 of culture, we observed generally decreasing cell confluency with decreased seeding density. To balance anticipated cell proliferation with efficient use of cells, the lowest seeding density was selected for subsequent studies.

### The gECM hydrogel wafers maintain adMSC viability and construct stability over long-term culture

In our study design, 1 human adMSC donor was seeded onto gECM hydrogel wafers made from 1 cartilage and 1 bone human tissue donor, with 3 total adMSC donors, 3 total cartilage tissue donors, and 3 total bone tissue donors (**Figure 5A**, **Table 2**). The adMSCs were seeded onto pre-hydrated gECM hydrogel wafers (70,000 cells/100µL wafer) and cultured for 21 days. Confocal imaging of the top surface showed high viability on Day 3 and Day 21, with increased cell count indicating proliferation (**Figure 5B**). Cells appear to localize and adhere to ECM particles by Day 21, evident by spindle-shaped cells on ECM particles. We observe a more cohesive network of cells on Day 21 in cartilage wafers compared to bone. Cross-sectional images reveal increased penetration of adMSCs into the depth of the wafers by Day 21, most notably in bone wafers. This may suggest some migration of cells through the porous internal network of the wafers. Minimal cells were observed on the bottom surface, suggesting that cells primarily reside near the top surface, but some cells did appear to migrate around the outside edge to the bottom surface in both cartilage and bone wafers.

**Figure 5.**
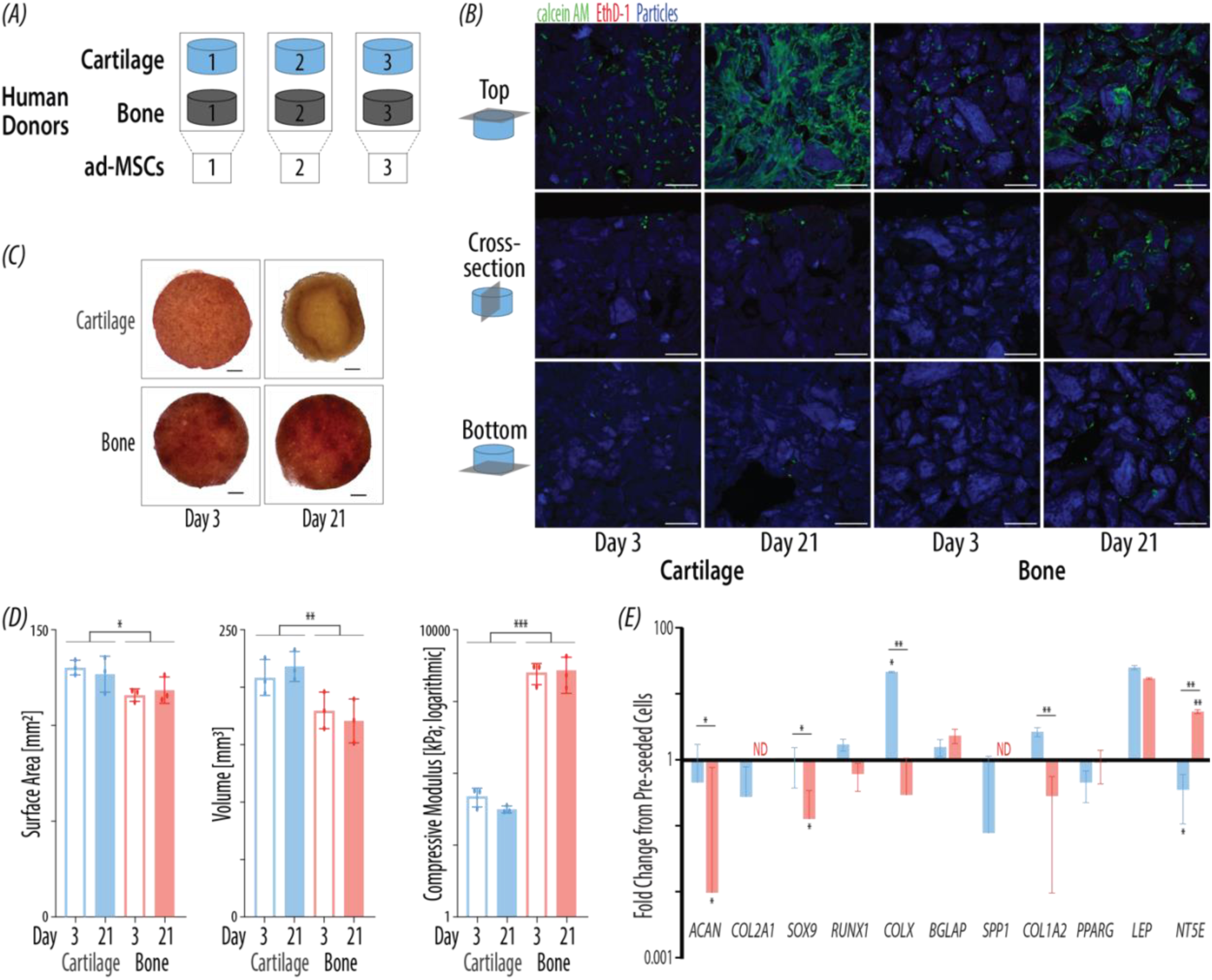
gECM hydrogel wafers maintain adMSC viability and construct stability over 21-day culture. (A) In our study design, one adMSC donor was seeded into one cartilage and one bone tissue donor gECM hydrogels (adMSC N = 3, gECM N = 3). (B) Confocal images of the top, cross-section, and bottom faces of cartilage and bone gECM hydrogels at Day 3 and 21. Green = LIVE, red = DEAD, blue = ECM particles autofluorescence. Scale bars = 250µm. (C) Representative macroscopic images of cartilage and bone gECM hydrogel wafers at Day 3 and 21. Scale bars = 1mm. (D) Surface area, volume, and compressive modulus at 20-30% strain (gECM N=3, adMSC N = 3) on Day 3 and 21. Error bars = standard deviation. (E) Quantitative PCR showing gene expression (gECM N = 3, adMSC N = 3), fold change to control adMSCs prior to seeding (N = 3 except LEP N=1). ND = not detectable. Error bars = SEM. For all plots, *p<0.05, **p<0.01, ***p<0.001.

We characterized geometry and mechanics of the gECM hydrogel wafers to assess their stability over long-term culture. Representative images (**Figure 5C**) demonstrate that wafers maintained overall integrity over culture time, with some macroscopic changes in cartilage wafers. Cartilage wafers underwent visible contraction, resulting in upward folding edges with a central concavity by Day 21. This observed geometric change is likely cell-mediated, driven by the adMSC network pulling particles together on the surface. Quantitatively (**Figure 5D**), there were no significant changes in wafer surface area or volume over the culture time. Cartilage wafers were significantly larger overall in both surface area and volume compared to bone, likely driven by wafer swelling patterns characterized earlier. Similarly, there were no significant changes in wafer bulk stiffness over the culture time, but bone exhibited overall significantly higher compressive modulus compared to cartilage. This mechanical stability was observed 4 regions of strain (0-10%, 10-20%, 20-30%, and 30-40%) (**Supplementary Figure 5**). Together, these results demonstrate that gECM hydrogel wafers remain structurally and mechanically stable over culture time, preserving relevant mechanical cues for cells.

Finally, we quantified gene expression of adMSCs on cartilage and bone gECM hydrogel wafers on Day 21, shown as fold change relative to control pre-seeded adMSCs (**Figure 5E**). Cartilage exhibited significantly higher expression of canonical chondrogenic markers, ACAN and SOX9 [18,19], compared to bone. Additionally, while COL2A1 was downregulated in cartilage wafers compared to control pre-seeded cells, it did not amplify at all for bone wafers, suggesting increased expression in cartilage compared to bone. However, cartilage also exhibited significantly higher expression of COLX and COL1A2 compared to bone, potentially suggesting hypertrophic behavior in cartilage [57]. We observe higher expression of BGLAP, a bone marker [58,59], in bone wafers compared to cartilage, but this difference is not statistically significant. We see some differences in expression relative to control pre-seeded cells, with bone significantly downregulating ACAN and SOX9 and upregulating NT5E, while cartilage significantly upregulated COLX. Only 1 control pre-seeded adMSC donor expressed LEP, a late adipogenic marker [60], so statistical analyses could not be completed. However, we did observe upregulation of LEP in both cartilage and bone wafers relative to control pre-seeded cells. While LEP can be expressed in adipogenesis [60], the paired downregulation of PPARG likely indicates osteogenesis instead [61]. NT5E was significantly downregulated in cartilage but upregulated in bone relative to control pre-seeded cells. Importantly, gene expression results were comparable to those quantified in adMSCs previously encapsulated in cartilage and bone gECM hydrogels (unpublished). This supports the ability of a 2.5D gECM hydrogel wafer substrate to drive similar responses as a 3D gECM hydrogel model. Taken together, these results suggest that cartilage and bone gECM wafers supported differential cell responses, with evidence of differentiation particularly in cartilage. While distinct chondrogenic and osteogenic phenotypes were inconclusive, 2.5D gECM hydrogel wafers drove similar cell activity as 3D gECM hydrogels.

### gECM hydrogel wafers can be assembled into microphysiological models with on-chip hydration and large biomolecule flow

Finally, we demonstrated incorporation of cartilage gECM hydrogel wafers into a proof-of-concept microphysiological on-chip model, with *in situ* hydration and large biomolecule flow. A single-channel PDMS chip was designed and fabricated containing a cartilage gECM hydrogel wafer (**Figure 6A**). Once assembled, DPBS was flowed into the tubing to hydrate the wafer (**Figure 6B**). Stereoscopic images demonstrate *in situ* swelling, with observed gaps between the dry wafer and channel walls, and no gaps between the swelled wafer and channel walls. We next introduced 250 kDa fluorescent Dextran to the chip, capturing confocal images before, during, and after Dextran flow (**Figure 6C**). This size range encompasses large biomolecules, representing a physiologically relevant upper bound for transport of molecules important to joint health and testing, including lubricin, growth factors, and therapeutic agents. Confocal images were captured 30µm up into the wafer, where we see optimal visualization of ECM particles. As Dextran is flowed into the chip, we see increased quantity of green staining between ECM particles, indicating flow through the wafer. We further captured images as Dextran was flushed out with DPBS, to demonstrate efficient removal of large biomolecules from the system. After flushing, minimal green staining is observed. Dextran was additionally flowed through the chip while confocal imaging 100µm into the wafer (**Supplementary Figure 6**). We observe less clear visualization of particles at this depth, but still see Dextran flow between particles, although quantity of Dextran appears lower at this depth compared to 30µm. Overall, we demonstrate successful incorporation of gECM hydrogel wafers into a proof-of-concept microphysiological model, with on-chip hydration and large biomolecule flow.

**Figure 6.**
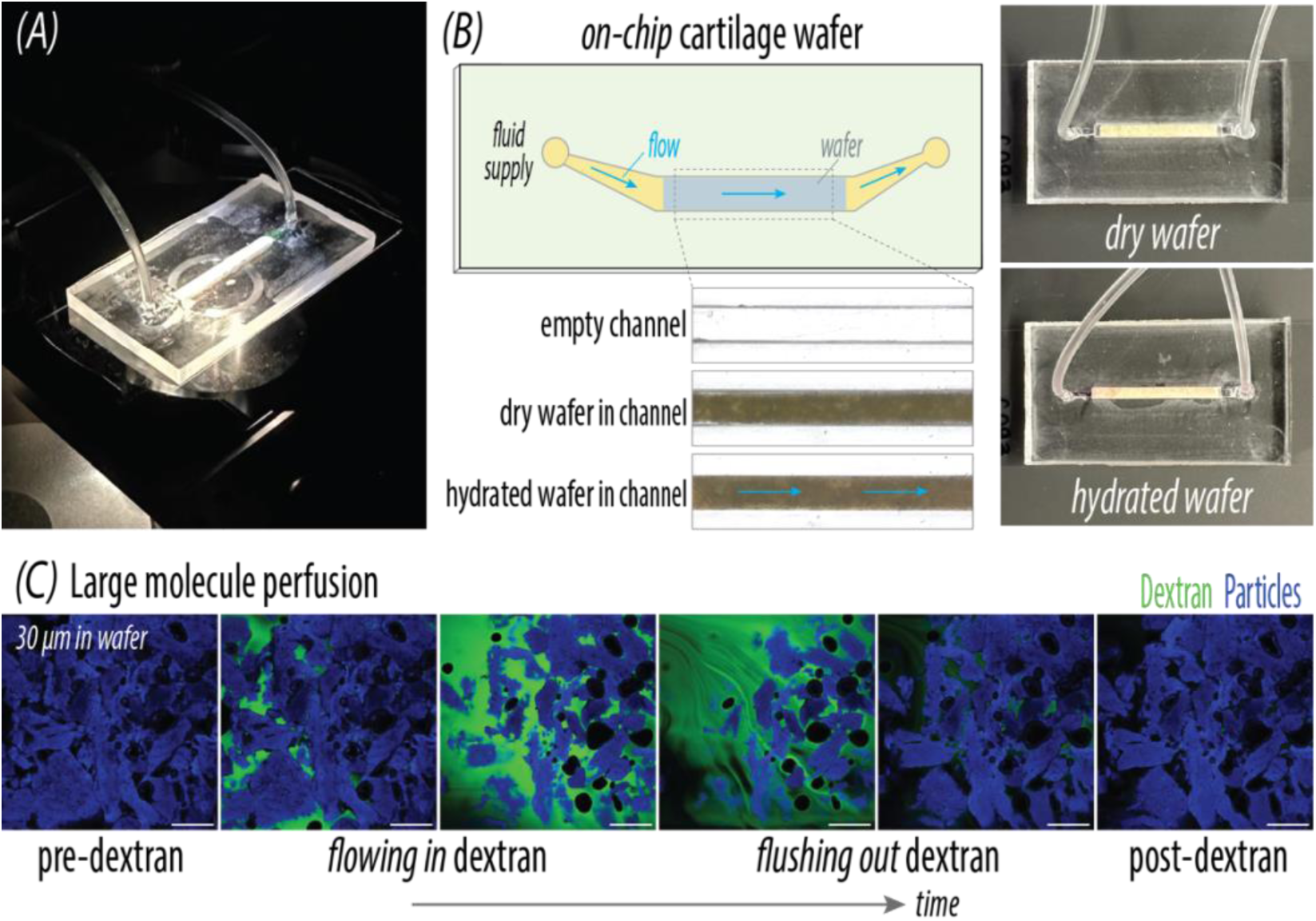
gECM hydrogel wafers can be assembled into microphysiological models with on-chip hydration and large biomolecule flow. (A) Picture of assembled chip on confocal microscope. (B) Stereoscopic images showing empty channel, dry wafer inside the chip channel, and hydrated wafer inside the chip channel (1hr hydration). Scale bar = 1mm. Pictures of assembled chip before and after *in situ* hydration (1hr hydration). (C) Confocal images taken 30µm into wafer before Dextran, while flowing in Dextran, while flushing out Dextran with DPBS, and after Dextran. Green = Dextran, Blue = particle autofluorescence. Scale bars = 250µm.

## DISCUSSION

In this study, we developed lyophilized granular extracellular matrix (gECM) hydrogel wafers as a 2.5D, shelf-stable, humanized substrate for *in vitro* modeling. These wafers were fabricated from human cartilage and bone-derived gECM hydrogels and retained physical cohesion upon rehydration, while maintaining key physical properties including swelling behavior, particle volume fraction, and bulk mechanics over a three-month period of room temperature storage. Importantly, we demonstrated that lyophilization transformed gECM hydrogels into a structurally distinct substrate characterized by increased porosity and defined surface topology, while preserving tissue-specific mechanical properties. We further established a simple and reproducible cell seeding strategy using pre-hydrated wafers, enabling sustained viability of human adMSCs over 21 days in culture. Cellular activity and gene expression trends were comparable to previously reported 3D gECM hydrogel systems, supporting the ability of this 2.5D platform to recapitulate key aspects of cell–material interactions without the need for encapsulation. Finally, we demonstrated integration of gECM wafers into a proof-of-concept microphysiological system, including *in situ* hydration and transport of large biomolecules, highlighting their potential for incorporation into complex *in vitro* models.

Lyophilization fundamentally altered the structure of gECM hydrogels, producing a wafer with more defined porosity. Compared to non-lyophilized hydrogels, which exhibited relatively smooth surfaces and a more continuous thiolated hyaluronic acid (tHA) phase, wafers displayed increased surface topography and more defined porosity in the inter-particle tHA. This increase in porosity is consistent with the formation of ice crystals during freezing, followed by sublimation during lyophilization, which leaves behind a pore network [46,52]. Despite this introduction of porosity, wafers exhibited increased particle volume fractions and corresponding increased compressive stiffness, likely drive by percolation, relative to their hydrogel counterparts established in previous work [manuscript in progress]. This suggests that lyophilization drives particles into a compacted configuration that persists upon rehydration. This increased particle compaction may explain the limited cell migration into the depth of the wafers following surface seeding, despite visually apparent porosity. Interestingly, cartilage wafers appeared to maintain this compacted state more effectively than bone wafers over storage time. This may be attributed to differences in particle composition and mechanics, where softer cartilage particles enriched in proteoglycans facilitate interparticle entanglement and stabilization of a jammed particle network [62]. In contrast, rigid mineralized bone particles may resist compaction and more readily relax toward their pre-lyophilized configuration. Together, these results indicate that gECM hydrogel wafers exhibit a structurally distinct architecture compared to gECM hydrogels, with more defined inter-particle tHA porosity but higher particle compaction.

A key advantage of the gECM wafer platform is the ability to seed cells directly onto the material without encapsulation, significantly simplifying experimental workflows. We established that pre-hydration of wafers prior to seeding was critical for maintaining cell viability, as human adMSCs seeded onto dry wafers exhibited increased penetration depth but substantially reduced viability. These findings are consistent with SEM observations, which showed that dry wafers possess sharper features and more pronounced microporosity, whereas hydrated wafers exhibit smoother surfaces with slightly reduced but still interconnected porosity. The sharper features in dry wafers may contribute to mechanical stress or shear-induced damage to cells. Limited immediate access to surrounding media during initial attachment may further compromise viability in cells seeded directly on dry wafers. In contrast, pre-hydrated wafers provide a less abrasive environment with increased media supply that supports cell attachment and survival, albeit with reduced penetration depth. While the current system supports robust viability, variability in seeding and partial cell loss during initial application remain limitations. Some cells were lost over the sides of the wafer prior to attachment, suggesting that future strategies to improve retention—such as adhesion of wafers to cover the entire surface of a cell culture dish—may improve reproducibility. Future work will also focus on enhancing cellular integration into the depth of the wafers through microstructural tuning. Techniques such as directional freezing, physical patterning, or lower particle packing densities could introduce aligned or channel-like pores that promote deeper infiltration [37,38,46,48–51]. Notably, directional freezing approaches in bulk hydrogels have been shown to enhance cellular integration, although often at the expense of mechanical properties [46]. The particle-based nature of gECM wafers may offer an opportunity to decouple these effects, enabling improved transport pathways without compromising stiffness.

Although gECM hydrogel wafers are implemented as 2.5D substrates, they retain key compositional, structural, and mechanical features associated with 3D systems. Previous work that established decellularization methods and processing of ECM particles demonstrated distinct proteomic profiles indicating tissue-specific composition [manuscript in progress]. Roughness analysis and SEM imaging confirmed the presence of tissue-specific surface topology and internal structure, while mechanical testing demonstrated preservation of tissue-dependent stiffness differences between cartilage and bone. Cells seeded onto wafers were observed to penetrate into the material to a limited depth, allowing interaction with both surface and internal microstructure. Relatively smoother isotropic surface topology has been shown to promote chondrogenesis, while relatively rougher directional surface topology has been shown to promote osteogenesis [63]. Our cartilage gECM wafers demonstrated smoother macro-scale surface topology compared to bone gECM wafers, but future work may optimize surface roughness at the micro-particle scale where cells adhere [64] and incorporate directional surface patterning for bone wafers. Importantly, gECM wafers supported sustained cell viability and induced gene expression trends comparable to those observed in fully encapsulated 3D gECM hydrogels. While clear lineage-specific differentiation was not observed, wafers were able to drive differential human adMSC responses between cartilage and bone substrates, suggesting that tissue-specific cues remain functionally active in the 2.5D format. These findings are consistent with prior work demonstrating that tissue type in 3D gECM hydrogels influence human adMSC behavior. Future work may focus on enhancing differentiation and cell response through material and biochemical optimization, including the use of tissue-specific hydrogel bases (e.g., hyaluronic acid for cartilage, collagen for bone), enhanced incorporation of growth factors within the media or material, or exploration of particle-only wafer systems that eliminate the hydrogel phase. Additionally, this platform may be particularly well-suited for maintaining the phenotype of already differentiated cells, as prior work with gECM hydrogels has demonstrated preservation of chondrocyte and dermal fibroblast phenotypes in tissue-matched hydrogels [manuscript in progress]. Importantly, the ability of gECM wafers to recapitulate key aspects of 3D systems while enabling direct cell seeding represents a significant advantage in usability. This platform effectively bridges the gap between traditional 2D systems, which lack physiological relevance, and 3D encapsulation approaches, which are often technically complex and less accessible for widespread use.

A defining feature of gECM wafers is their ability to be stored dry at room temperature while maintaining structural and functional properties over time. Over a three-month period, we observed no significant changes in swelling behavior, particle network structure, or mechanical properties, suggesting that lyophilization stabilizes the material without compromising performance. This shelf stability addresses a major limitation of conventional hydrogel systems, which typically require immediate use following fabrication and often rely on refrigerated or frozen storage. In contrast, dry porous biomaterials such as collagen or gelatin sponges are known to exhibit multi-year shelf lives under appropriate conditions [39,40,53], suggesting that gECM hydrogel wafers may similarly achieve extended stability beyond the timeframes evaluated in this study. Importantly, shelf stability enables new modes of integration into microphysiological systems. We demonstrated that dry gECM hydrogel wafers can be incorporated into a channel on a microfluidic device, with *in situ* swelling that enables wafer conformity to the channel geometry. This pre-assembly approach contrasts with many existing systems that require biomaterials to be introduced in a hydrated or cell-laden state, limiting flexibility in device fabrication and workflow design. In addition, we demonstrated transport of large biomolecules (250 kDa Dextran) through the wafer structure, indicating that the porous network supports flow of physiologically relevant molecules such as growth factors, serum proteins, and biologic therapeutics. Notably, the on-chip cartilage gECM hydrogel wafer was only hydrated for 30 mins before flowing Dextran. The majority of swelling for cartilage gECM hydrogel wafers happens in 30 mins, although 24 hours is required to reach a full plateau. Future studies should assess large molecule flow after the wafer has been fully hydrated to mimic introduction of large biomolecules at later time points during on-chip culture. While transport of Dextran appeared reduced at greater depths within the wafer, this may be influenced by short exposure duration, where longer duration might enable diffusion of molecules deeper into wafers. Importantly, as cells were primarily localized near the surface, the observed transport is already sufficient to support relevant cell–molecule interactions, though improved cellular infiltration would likely enhance transport of biomolecules throughout the construct as well.

Overall, gECM hydrogel wafers represent a versatile platform that combines biomimicry, ease of use, and shelf stability. By bridging the usability of 2D substrates with the biological relevance of 3D ECM-based materials, this 2.5D approach enables more accessible implementation of physiologically relevant *in vitro* models. Future work will focus on tuning microstructure to improve cellular integration and retention, including strategies such as directional freezing and physical patterning. Expansion of this technology to a variety of tissue types will further enhance the broad utility of this platform. Together, these efforts position gECM wafers as a promising tool for advancing microphysiological modeling and improving translational outcomes in tissue engineering and drug discovery.

## MATERIALS AND METHODS

### Tissue Particle Processing

Human articular cartilage (N=6 human donors, **Table 1**) and cancellous bone (N=10 human donors, **Table 1**) from the knee joint were sourced from AlloSource, a human donor tissue bank, and separated by donor. Cartilage was cut into ∼4mm^3^ pieces and decellularized using a viral inactivation method [56]: orbital agitation at RT in 0.2M HCl for 2hrs, 1.0M NaOH for 2hrs, and rinsing in 0.02M HCl and water/DPBS until neutral (pH=7.5). Bone was pulverized and cleaned following the AlloSource AlloTrue cleanse process, including rinses in hydrogen peroxide, isopropyl alcohol, and an antibiotic solution. Cartilage and bone was then frozen, lyophilized, pulverized in a stainless-steel milling jar (Tissue Lyser III, Qiagen), and size sorted to <250 µm diameter using a sieve. Bone then underwent an additional de-fat process, including an 8hr orbital agitation in ethanol (50%, 75%, 95%, 100% for 2hrs each) [65], and overnight orbital agitation in DPBS. It was then lyophilized and sorted to include particles 40-250µm diameter.

**Table 1.**
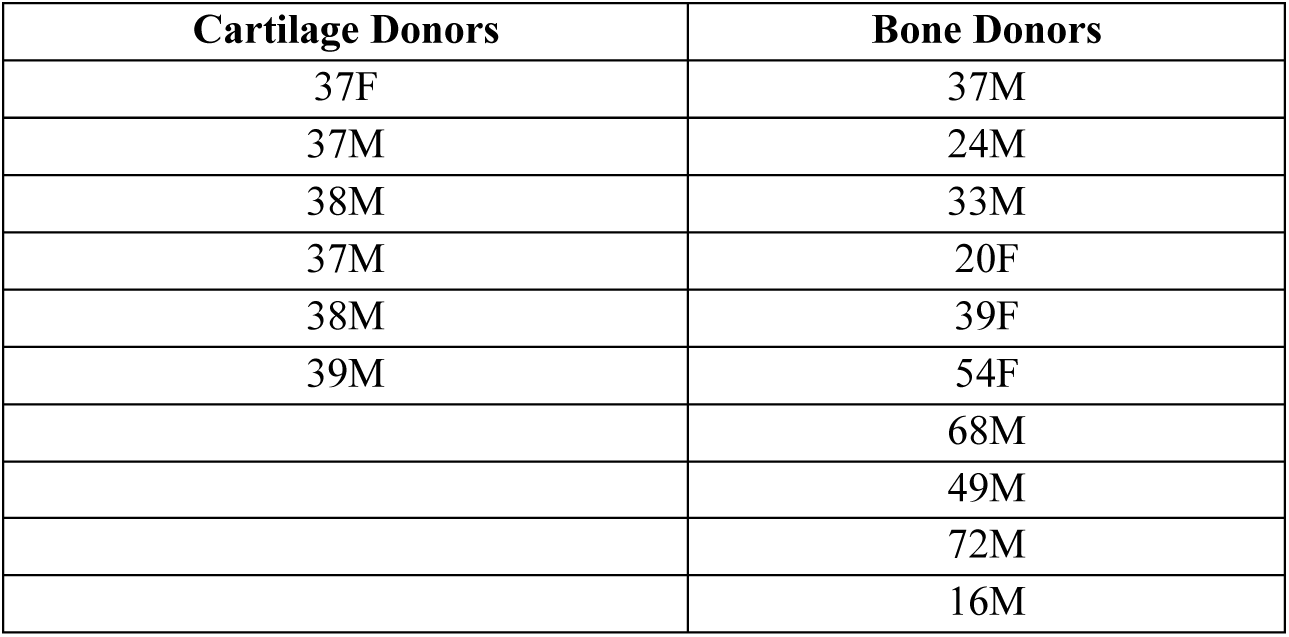
Age and sex (Male, Female) of human cartilage/bone donors used for wafer characterization.

### gECM Hydrogel Wafer Fabrication

#### Hyaluronic Acid Functionalization

Glucoronate carboxyl groups on hyaluronic acid (Lifecore Biomedical, HA-100K-5) were replaced with thiol groups following previously established protocols to produce thiolated hyaluronic acid (tHA) [23], with purification via tangential flow filtration. The substitution rate was confirmed to be 18%–23% (thiolated mmols/unthiolated mmols) using a standard Ellman’s assay (Ellman’s solution, ThermoFisher).

#### gECM Hydrogel Mixing

The gECM hydrogel formulations consisted of 10 mg/mL tHA packed with 0.2 g/mL (cartilage) [23] or 1 g/mL (bone) lyophilized particles. The gECM hydrogels were molded into cylindrical constructs (1.5 mm height x 5 mm diameter) and incubated for 45 min (cartilage) or overnight (∼16hours, bone) at 37°C.

#### gECM Hydrogel Wafer Fabrication

Polymerized gECM hydrogel constructs were isotropically frozen at -20C for ∼3 hrs and then lyophilized for ∼24 hrs to form gECM hydrogel wafers.

### gECM Hydrogel Wafer Characterization

#### Swelling

The gECM hydrogel wafers were incubated in DPBS at room temperature for 1 week. Mass and volume were measured first in the dry wafer form (0 hr), and then at 6 timepoints during incubation in DPBS (30 mins, 1 hr, overnight ∼16hrs, 24 hrs, 48 hrs, 1 wk). Volume was obtained by measuring height with calipers and calculating surface area from thresholded (Image J) stereoscopic images (Leica).

#### Particle Volume Fraction

Tissue particles in fully-swelled gECM hydrogel wafers were stained with Ghost Dye 710 (Cytek Biosciences) and imaged via confocal microscopy (Nikon A1R, 20x objective, NA=0.75). We obtained three-dimensional 5µm-step z-stack images at 3 different locations per sample. Using a custom MATLAB code, we selected, thresholded, and averaged the area over 15µm depth to calculate particle volume to total construct volume.

#### Bulk Compression

Fully-swelled gECM hydrogel wafers were compressed (unconfined) to 40% strain at a quasi-static 0.1%/s strain rate (MCR 702, Anton Paar). Compressive modulus was calculated between 0-10%, 10-20%, 20-30%, and 30-40% strain [23].

### gECM Hydrogel Wafer Shelf-Life Study

Fresh gECM wafers were fabricated and immediately characterized. gECM wafers for shelf-life studies were fabricated, stored in moisture-resistant packaging at room temperature, and then characterized at 1 month or 3 months.

### gECM Hydrogel Microstructure and Topography

#### Microstructure

Microstructure was visualized with scanning electron microscopy (SEM). Cartilage and bone gECM hydrogels (NO lyophilization), gECM hydrogel wafers (lyophilized, dry form), and hydrated gECM hydrogel wafers (lyophilized, then hydrated in PBS) were imaged. gECM hydrogels and hydrated gECM hydrogel wafers were first equilibrated in DPBS for 24-48hrs, followed by a graded ethanol treatment (20% ethanol 2hrs, 40% ethanol 2hrs, 60% ethanol overnight, 80% ethanol 2hrs, 100% ethanol overnight) and critical point drying (Leica EM CPD300). All samples were fractured to expose internal structure, sputter coated with 10nm platinum (Leica EM ACE600), and SEM imaged (Hitachi High-Tech SU3500) at an accelerating voltage of 10 kV at multiple magnifications (top and cross section).

#### Surface Roughness

Surface topology of both dry and hydrated (in DPBS for 24-48hrs) cartilage and bone gECM hydrogel wafers were quantified using a laser scanning microscope with a 20x objective (Keyence VK-X1000 Series). At 3 areas per sample, arithmetic mean height, root mean square height, surface skewness, and surface kurtosis were extracted from 3D surface reconstructions using the Keyence analysis software.

#### Optical Coherence Tomography

Optical coherence tomography (OCT) images were taken (Thorlabs OCT-TEL220C1, A-scan rate: 5.5kHz) of cartilage and bone gECM hydrogel wafers in dry form (T0), immediately after adding PBS (T1), and after 24hrs in PBS. Height was measured at various points along cross-sectional 2D views and averaged to obtain a mean height value at each time point. Isometric 3D views were used to visualize surface topography.

### adMSC Culture

#### Expansion

Cryopreserved, human adipose-derived MSC’s (N=3 human donors, **Table 2**) were sourced from AlloSource, thawed, and expanded to P4 in StemMACS MSC Expansion medium (Miltenyi Biotec).

**Table 2.** Age and sex (Male, Female) of human adMSC and human cartilage/bone donors, grouped for cell studies.

| **adMSC Donor** | **Cartilage Donor** | **Bone Donor** |
| --- | --- | --- |
| 28F | 37F | 37M |
| 23M | 38M | 16M |
| 21M | 37M | 72M |

#### Seeding Method Pilot

One set of cartilage (N=1) and bone (N=1) gECM hydrogel wafers (1.5 mm height x 10 mm diameter) were placed in non-tissue-culture-treated plates and pre-hydrated in culture medium for 24-48hrs, and then media was removed. Another set of gECM wafers was placed in non-tissue-culture-treated plates in dry form. P4 adMSCs suspended in culture medium were seeded on top of gECM hydrogel wafers (500,000 cells / 100µL wafer). Cartilage or bone-specific media was added to cell-laden gECM hydrogel wafers 2hrs after seeding. On Day 3, we assessed cell viability (Calcein AM, Ethidium Homodimer-1; ThermoFisher Invitrogen) via confocal microscopy (Nikon A1R, 10x objective, NA=0.45).

#### Seeding Density Pilot

Different seeding densities (250,000 cells, 125,000 cells, or 70,000 cells / 100µL wafer) were tested on cartilage and bone pre-hydrated (24-48hrs in culture medium) gECM hydrogels. Cartilage or bone-specific media was added to cell-laden gECM hydrogel wafers 2hrs after seeding. On Day 3, we assessed cell viability (Calcein AM, Ethidium Homodimer-1; ThermoFisher Invitrogen) via confocal microscopy (Nikon A1R, 10x objective, NA=0.45)..

#### Seeding and Culture

Cartilage (N=3) and bone (N=3) gECM hydrogel wafers were placed in non-tissue-culture-treated plates and pre-hydrated in cartilage or bone culture medium for 24-48hrs. Medium was removed, and adMSCs suspended in culture medium were then placed on top of the hydrated gECM hydrogel wafers (70,000 cells / 100µL wafer). Media was added to cell-laden gECM hydrogel wafers 2hrs after seeding. Cartilage gECM hydrogel wafers were cultured in cartilage media (Dulbecco’s Modified Eagle Medium DMEM supplemented with 2% FBS, 1% ITS+Premix, 1X sodium pyruvate, 1X Pen/Strep, 50µg/mL ascorbic acid, 100nM dexamethasone, and 2.5ng/mL TGFβ3) and bone gECM hydrogel wafers were cultured in bone media (Dulbecco’s Modified Eagle Medium DMEM supplemented with 2% FBS, 1% ITS+Premix, 1X sodium pyruvate, 1X Pen/Strep, 50µg/mL ascorbic acid, 10mM β-glycerophosphate, and 100nM dexamethasone). Cell-laden gECM hydrogel wafers were cultured for 21 days, changing medium every 3-4 days. Cell-gECM constructs were transferred to new non-tissue-culture-treated plates on Day 3.

#### Viability

On Day 3 and 21, we assessed cell viability (Calcein AM, Ethidium Homodimer-1; ThermoFisher Invitrogen) on the top, bottom, and cross-section of gECM hydrogel wafers via confocal microscopy (Nikon A1R, 10x objective, NA=0.45). Particles were visualized via autofluorescence in the DAPI channel.

#### Construct Geometry and Bulk Stiffness

Construct geometry and bulk stiffness were quantified on Day 3 and 21. Surface area was calculated by thresholding (Image J) stereoscopic images of cell constructs. Height was measured using a BOSE ElectroForce 5500 system and used in combination with surface area to calculate volume. Cell constructs were compressed (unconfined) to 40% strain at a quasi-static 0.1%/s strain rate (BOSE ElectroForce 5500). Compressive modulus was calculated between 0-10%, 10-20%, 20-30%, and 30-40% strain [23].

#### Gene Expression

Gene expression was assessed on Day 21 through quantitative real-time PCR (CFX96 Touch). Samples were lysed in Qiazol and RNA was isolated using a Direct-zol RNA Miniprep Kit (Zymo). RNA was reverse transcribed into cDNA using iScriptTM (BioRad). Real-time quantitative PCR (RT-qPCR) was performed with SsoAdvancedTM Universal SYBR® (BioRad) in a CFX96 thermocycler (BioRad). We targeted known genes to assess chondrogenic phenotype (ACAN, COL2A1, SOX9, RUNX1, COLX), osteogenic phenotype (BGLAP, SPP1, COL1A2), adipogenic phenotype (PPARG, LEP), and stemness (NT5E). All measurements were normalized to the housekeeping gene (GAPDH) and fold changes were measured from gene expression of adMSCs before seeding into gECM biomaterials.

### Microphysiological Model Demonstration

#### Chip Assembly and Hydration

A polycarbonate mold for a single-channel chip was designed and machined. Sylgard 184 polydimethylsiloxane (PDMS) was poured into the mold and cured to form the chip. Tubing was connected to the inlet and outlet of the PDMS chip. Cartilage gECM hydrogel wafers were fabricated in the same geometry as the chip channel. The gECM hydrogel wafer was placed into the chip channel, and the PDMS chip surface was plasma treated and bonded to a plasma-treated glass coverslip. DPBS was flowed into the tubing until the channel was filled and left for 1hr. Stereoscopic images (Leica) were taken of the chip channel and cartilage gECM hydrogel wafer before assembly, after assembly (dry wafer), and after hydration.

#### Dextran Flow

A cartilage gECM hydrogel wafer-on-chip was pre-hydrated with PBS. Dextran fluorescein (MW 250kDa, 2.5mg/mL in DI water) was flowed through the chip and then flushed out again with DPBS over a time period of 6 minutes. Confocal images were captured before, during, and after Dextran flow (Nikon A1R, 10x objective, NA=0.45). Particles were visualized via autofluorescence in the DAPI channel.

### Statistical Analysis

Mixed-model analyses of variance (ANOVAs) were performed on linear mixed models with tissue type and treatment (e.g. time, swell state, etc) as co-factors, donor (tissue donor and/or cell donor) as random effects, and the resulting measurement variable as the response (swelling volume, swelling mass, volume fraction, compressive modulus, surface area, volume, etc). Statistical analyses for gene expression were performed on ΔCq values. All residuals were checked, and if a non-normal distribution was identified, the data was transformed for statistical analysis. Post hoc multiple comparisons were performed using Tukey’s honestly significant difference (HSD) correction. Significance was defined as p<0.05.

## SUPPLEMENTARY FIGURES

**Supplemental Figure 1.**
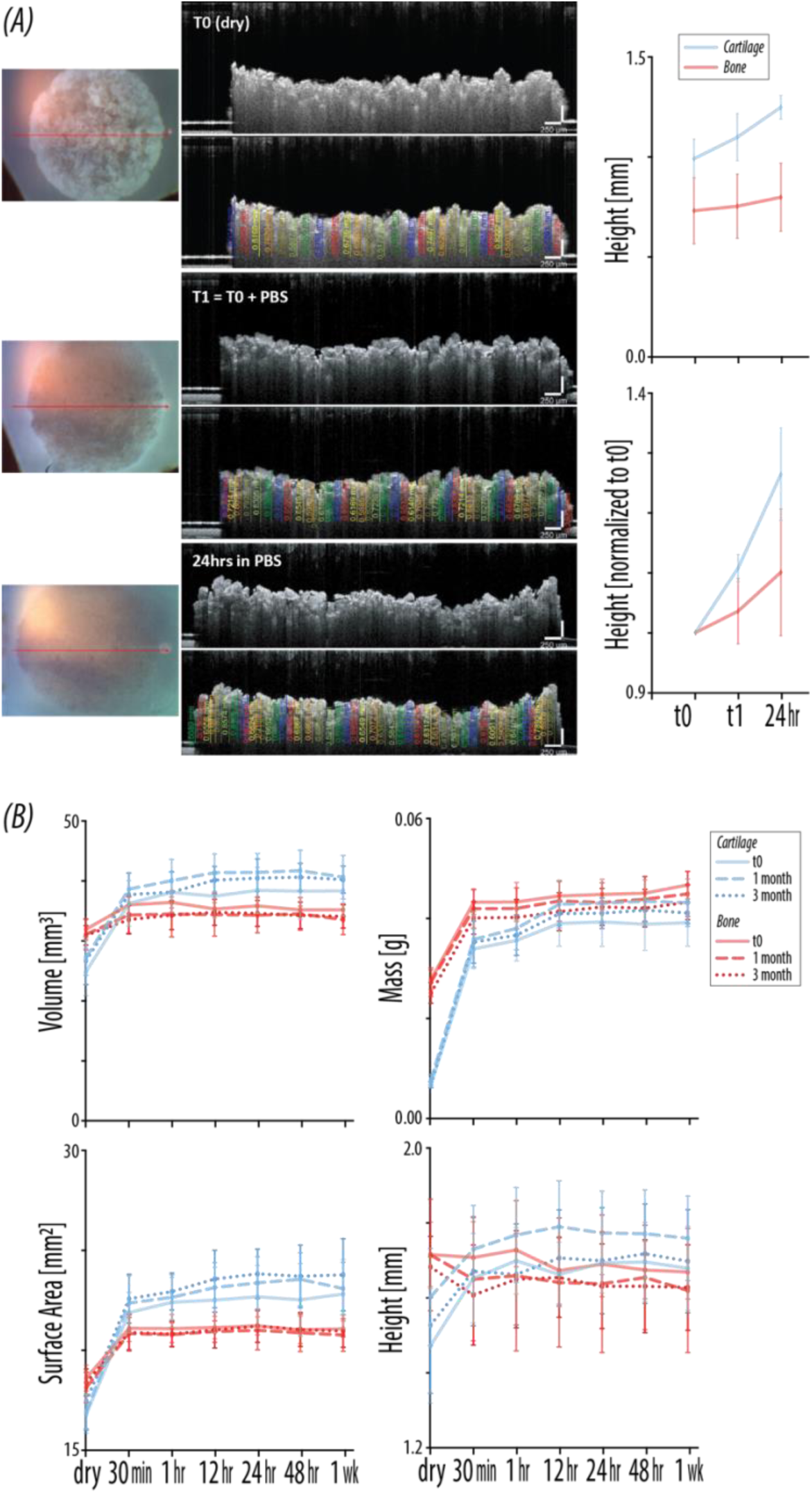
Additional swelling data. (A) Height (raw and normalized to t0) measured from cross-sectional optical coherence tomography (OCT) images at t0 (dry), t1 (PBS added and immediately imaged), and at 24hrs in PBS (N = 3-4). Height was measured at various points along the cross-section and averaged to obtain a mean height value. (B) Raw values for mass, volume, surface area, and height for gECM hydrogel wafers over 1 week swelling in PBS (N = 6). For all plots, error bars = standard deviation.

**Supplemental Figure 2.**
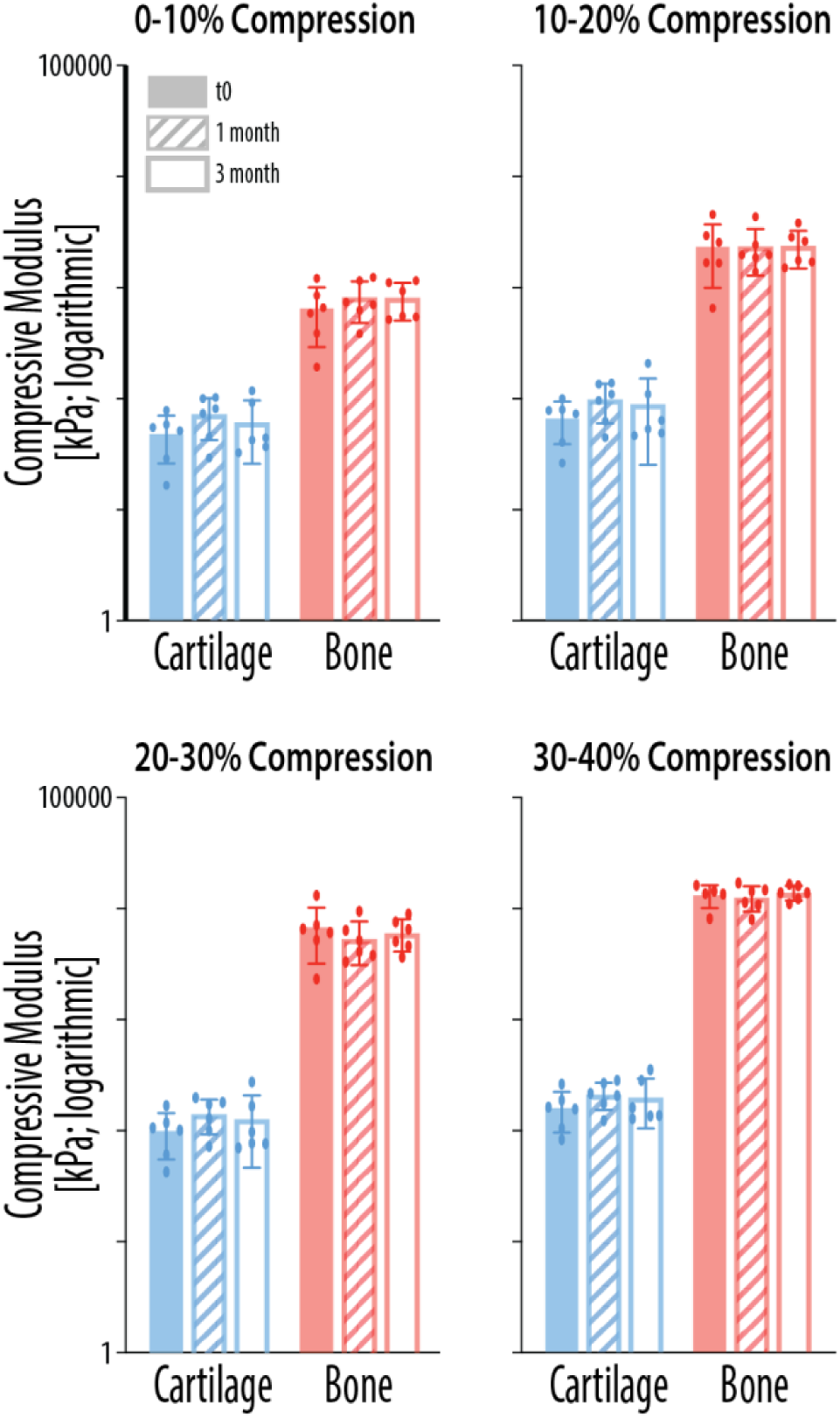
gECM hydrogel wafer compressive modulus calculated for various strain windows. Compressive modulus calculated at 0-10%, 10-20%, 20-30%, and 30-40% strain for gECM hydrogel wafers (N = 6). Error bars = standard deviation.

**Supplemental Figure 3.**
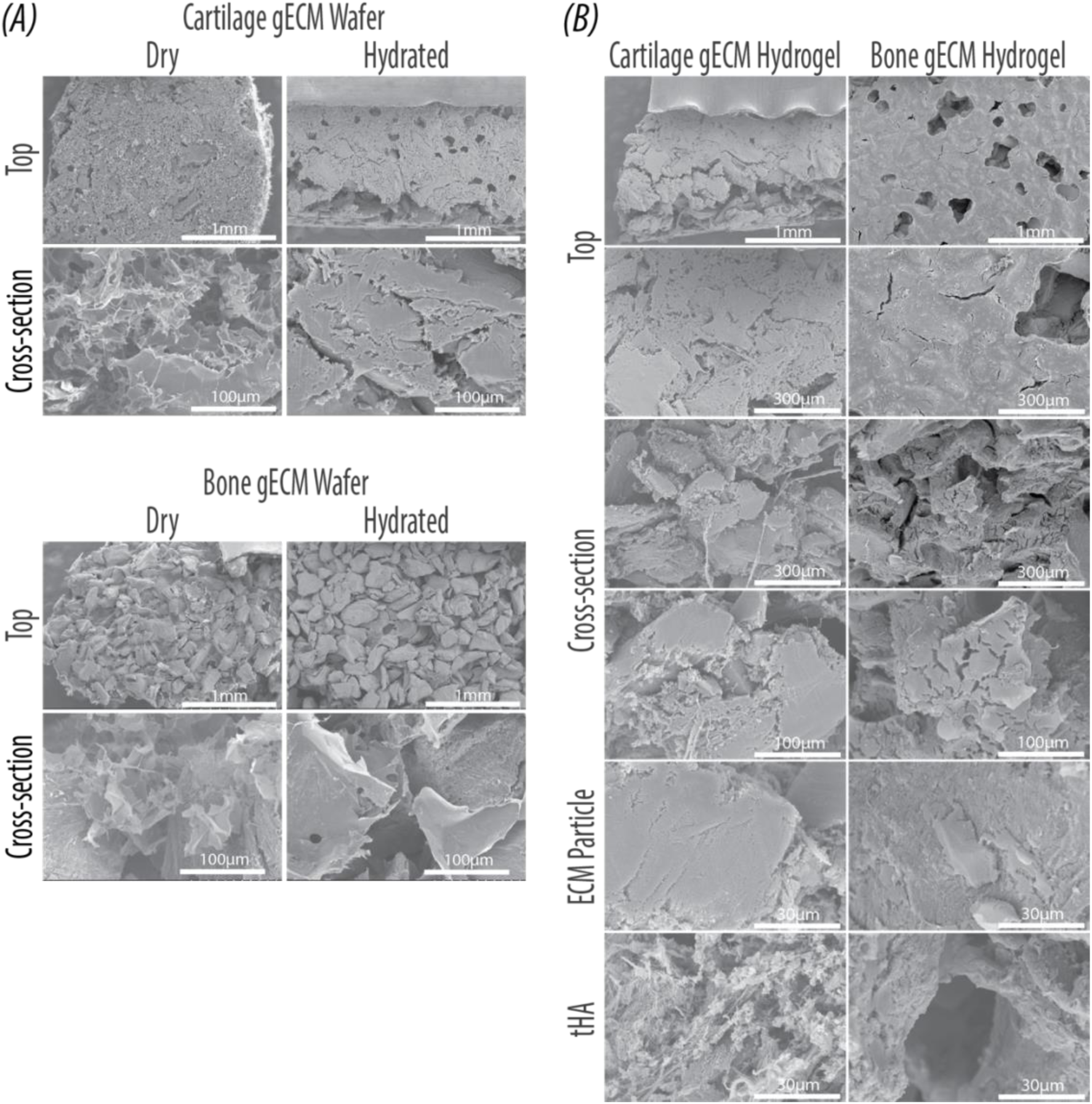
Scanning electron microscopy of gECM hydrogels. (A) Representative SEM images of cartilage and bone gECM hydrogel wafers in dry and hydrated (DPBS >24hrs) form. Image of the top are at 50X magnification (scale bars = 1mm) and cross-section are at 150X magnification (scale bars = 100µm). (B) Representative SEM images of cartilage and bone gECM hydrogels (NOT lyophilized). Images of the top are at 50X (scale bars = 1mm) and 450X magnification (scale bars = 300µm). Cross-section images are at 450X (scale bars = 300µm) and 150X magnification (scale bars = 100µm). Image of the particle and tHA are at 1500X magnification (scale bars = 30µm).

**Supplemental Figure 4.**
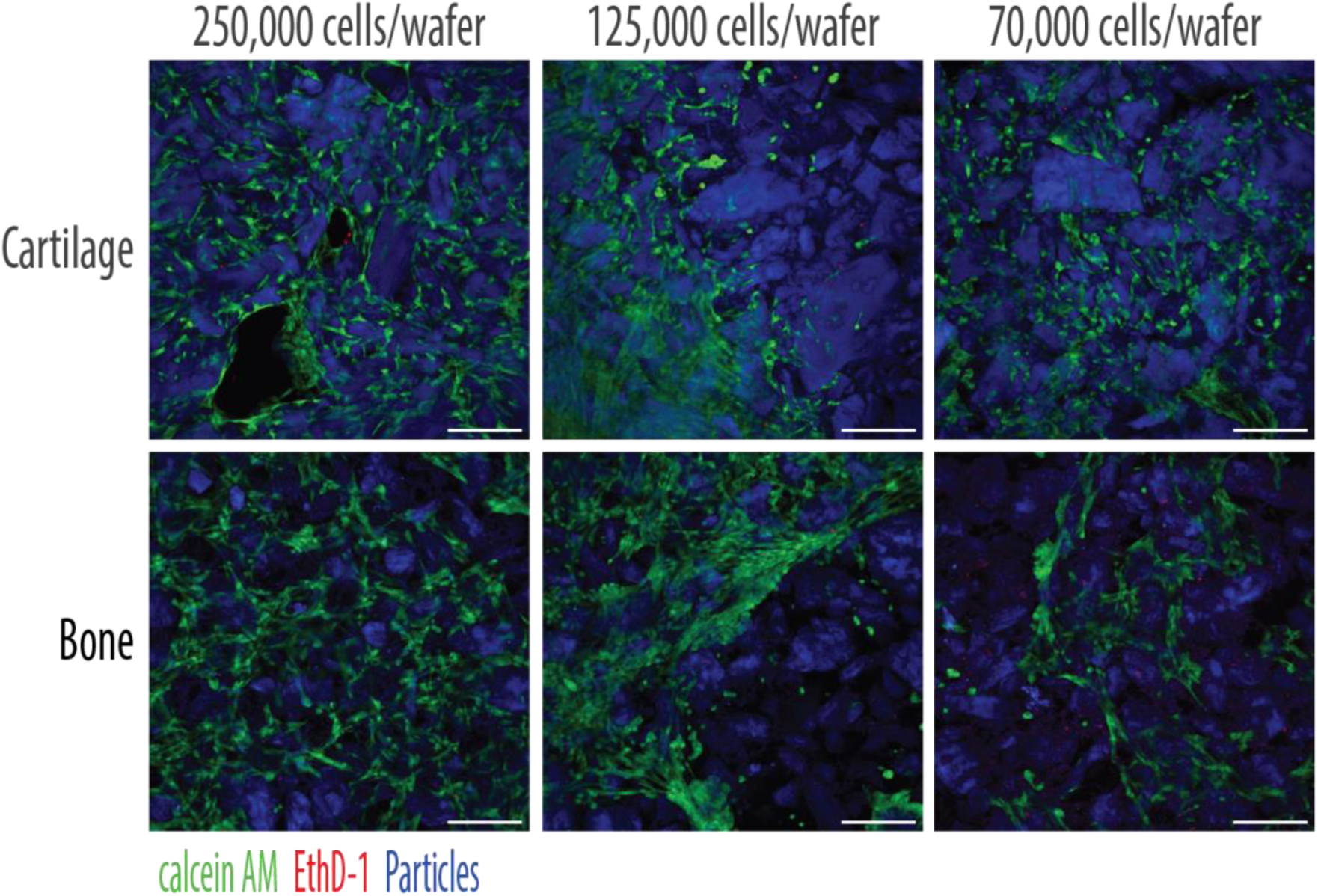
Cell seeding density. Ad-MSC’s seeded at 250,000 cells, 125,000 cells, and 70,000 cells per 100µm wafer. Confocal images of the top face of pre-hydrated cartilage and bone gECM hydrogel wafers at Day 3. For all images, green = LIVE, red = DEAD, blue = ECM particles autofluorescence, Scale bars = 250µm.

**Supplemental Figure 5.**
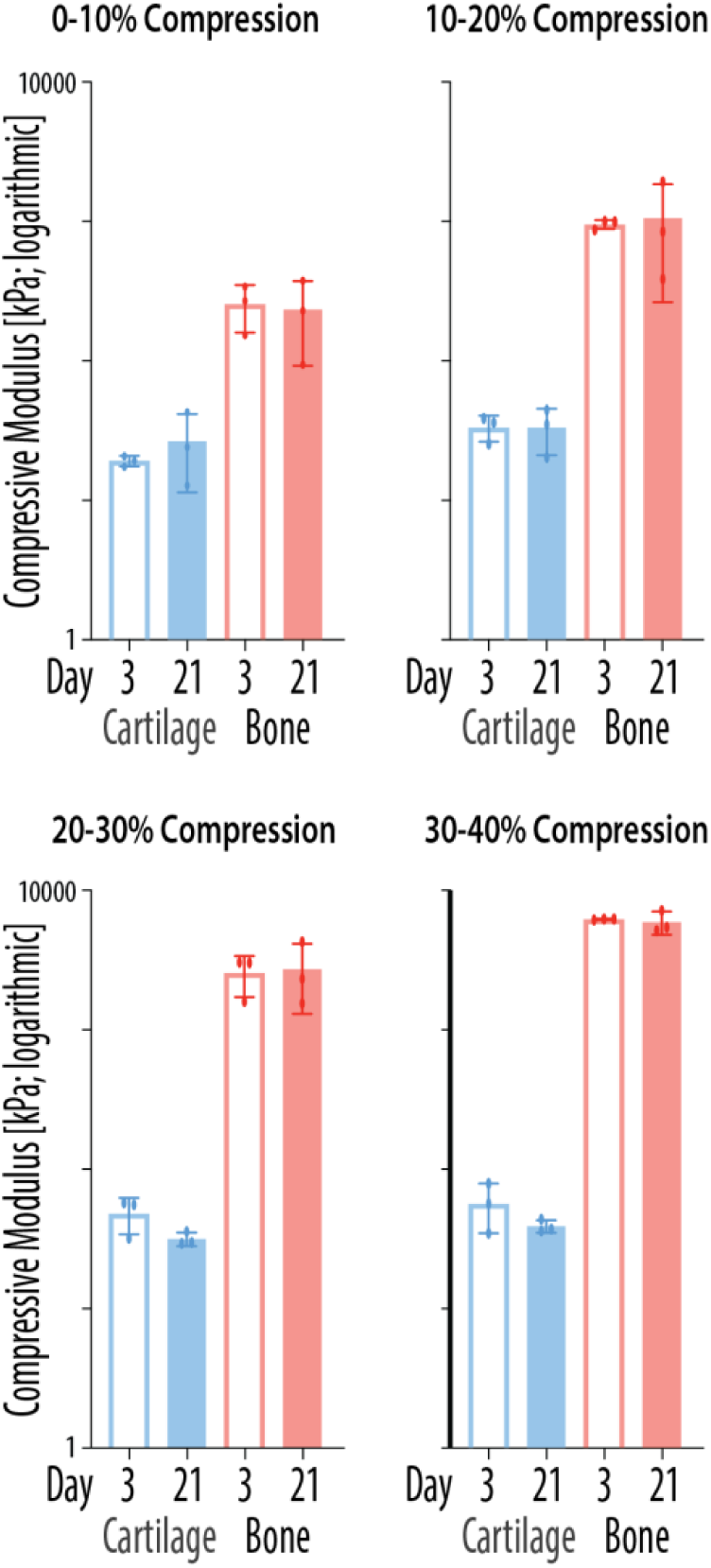
adMSC-laden gECM hydrogel wafer compressive modulus calculated for various strain windows. Compressive modulus calculated at 0-10%, 10-20%, 20-30%, and 30-40% strain for all tissue gECM hydrogel wafers (gECM N = 3, adMSC N = 3) on Day 3 and 21. Error bars = standard deviation.

**Supplemental Figure 6.**
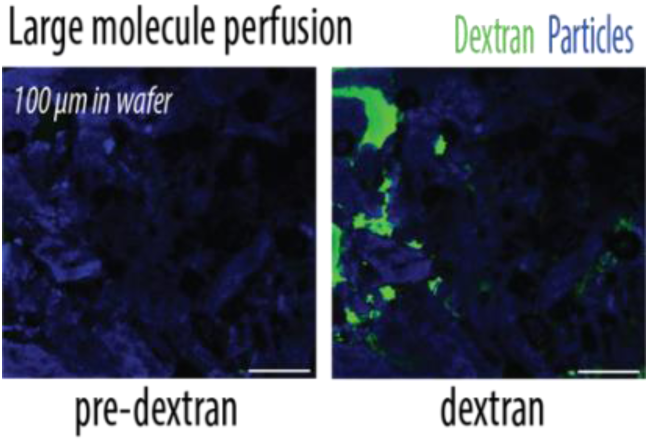
Dextran flow imaged 100µm into wafer. Confocal images taken 100µm into wafer before and while Dextran is flowed in. Green = Dextran, Blue = particle autofluorescence. Scale bars = 250µm.

## Acknowledgments

The authors acknowledge AlloSource for providing human tissues and human adMSCs. Critical point drying, sputter coating, and SEM imaging was performed at COSINC-CHR, CU Boulder (RRID:SCR_018985).

## Author contributions

Conceptualization: JOH, JEB, CPN

Methodology: JOH, SAB, SES, SG, JEB, MF, VA, SA, CPN

Visualization: CPN, JOH

Funding acquisition: CPN, JOH, SES

Supervision: CPN, SES

Writing – original draft: JOH

Writing – review & editing: All authors

